# Exploring the Landscape of Ectodomain Shedding by Quantitative Protein Terminomics

**DOI:** 10.1101/2020.09.23.310102

**Authors:** Kazuya Tsumagari, Chih-Hsiang Chang, Yasushi Ishihama

**Author notes:** Correspondence, /.

## Abstract

Ectodomain shedding is a proteolytic process that regulates the levels and functions of membrane proteins. Dysregulated shedding is linked to severe diseases, including cancer and Alzheimer’s disease. However, the exact cleavage sites of shedding substrates remain largely unknown. Here, we explore the landscape of ectodomain shedding by generating large-scale, cell-type-specific maps of shedding cleavage sites. By means of N- and C-terminal peptide enrichment and quantitative mass spectrometry, we quantified protein termini in the culture media of 10 human cell lines and identified 411 cleavage sites on the ectodomain of 132 membrane proteins whose proteolytic terminal fragments are downregulated in the presence of a broad-spectrum metalloprotease inhibitor. A major fraction of the presented cleavage sites was identified in a cell-type-specific manner, and mapped onto receptors, cell adhesion molecules, and protein kinases and phosphatases. We confidently identified 86 cleavage sites as metalloprotease substrates by means of knowledge-based scoring.

## Introduction

Membrane proteins have critical physiological roles, and their abundance and functions are tightly controlled through multiple mechanisms, including ectodomain shedding (shedding), which is a form of limited proteolysis that liberates the extracellular domain of membrane proteins (Huovila et al., 2005; Lichtenthaler et al., 2018; Weber and Saftig, 2012). Shedding contributes to cellular interactions with the environment by releasing active cytokines, growth factors or other mediators from their membrane-bound precursors, or conversely, by reducing the levels of receptors and adhesion proteins at the cell surface (Huovila et al., 2005; Weber and Saftig, 2012). Shedding often triggers further proteolysis within the transmembrane domain – known as regulated intramembrane proteolysis (RIP) – thereby releasing the intracellular domain into the cytoplasmic region. These processes enable bi-directional signal transduction (Beard et al., 2019; Reiss and Saftig, 2009; Weber and Saftig, 2012).

A protease involved in shedding is specifically referred to as a sheddase. Distinct sheddases may cleave the same substrate at different sites, generating multiple proteoforms that may exhibit different biological functions and activities (Niedermaier and Huesgen, 2019). For instance, the difference of cleavage sites of amyloid precursor protein (APP) by a disintegrin and metalloprotease 10 (ADAM10) and by beta-site APP cleaving enzyme 1 (BACE1) is critical for the pathogenesis of Alzheimer’s disease (Chow et al., 2010). Therefore, it is important to identify cleavage sites in order to achieve a precise understanding of the physiological roles of shedding.

Discovery and analysis of large numbers of proteoforms is the exclusive domain of liquid chromatography/tandem mass spectrometry (LC/MS/MS)-based proteomics (Aebersold et al., 2018; Olsen and Mann, 2013). For the identification of shedding substrates, it is preferable to investigate the proteins secreted into the cell culture media, rather than those remaining in the membrane, because the secretome is much less complex than the membrane proteome. Additionally, the fragments remaining in the membrane are often unstable due to further proteolytic events such as RIP and lysosomal degradation (Güner and Lichtenthaler, 2020; Merilahti and Elenius, 2019). By using established hydrazide chemistry, 18 metalloprotease substrates in the culture media were identified (Tsumagari et al., 2017); most of the membrane proteins are glycosylated (Apweiler et al., 1999). However, this approach did not lead to precise identification of the cleavage sites. In fact, knowledge of the cleavage sites of shedding substrates is limited, despite its importance. In standard shotgun proteomics, only peptides delimited by the specificity of the employed digestive enzymes are considered. Moreover, there is difficulty in consistently identifying proteolytic termini generated by shedding in semi-specific searches, since peptides may result from specific cleavage only at one terminus, while the other terminus may be generated by non-specific cleavage (Niedermaier and Huesgen, 2019). Thus, terminal peptide enrichment from secreted proteins coupled with semi-specific search should be employed in order to efficiently identify shedding cleavage sites.

Prudova *et al*. employed the terminal amine isotopic labeling of substrates (TAILS) method (Kleifeld et al., 2010) for N-terminal peptide enrichment and identified 201 and 19 cleavage products formed by recombinant MMP-2 and MMP-9, respectively (Prudova et al., 2010). However, there are still obstacles to the large-scale detection of cleavage sites generated by endogenous sheddases in living cells. First, terminal peptide enrichment strategies generally consist of multiple steps including some chemical reactions, and therefore relatively large amounts of samples are needed. Second, the protein amounts in the culture media are much smaller than those obtained from cell lysates, and thus it is hard to prepare sufficiently large samples for terminal peptide enrichment. Finally, C-terminomics specifically remains a challenging task, because the enrichment efficiency is greatly inferior to that of the N-terminal counterpart due to the difficulty in chemically modifying the carboxy group (Niedermaier and Huesgen, 2019).

Recently, our group developed a simple and rapid methodology for N-terminal peptide enrichment, in which N-terminal peptides of TrypN digests are isolated by strong cation exchange (SCX) chromatography at low pH (Chang et al., 2020). Earlier reports have demonstrated that C-terminal peptides generated by trypsin digestion are eluted first in SCX, together with acetylated N-terminal peptides (Alpert et al., 2010; Gauci et al., 2009; Helbig et al., 2010), and we discovered here this feature enables the identification and quantification of C-terminal peptides on a comparable scale to that of the N-terminal counterparts with the same amount of input (Figure S1A, B). Here, we applied these sensitive terminomics methodologies to conduct the first large-scale study of shedding cleavage sites targeted by endogenous metalloproteases, which have emerged as the major sheddase family. We quantitatively identify putative metalloprotease-regulated ectodomain shedding cleavage sites based on the results of broad-spectrum metalloprotease inhibitor treatment, and provide an overview of their positional and functional landscape.

## Results

### Sensitive terminomics workflow enables efficient identification and reproducible quantification of N- and C-termini in the secretome

To achieve large-scale detection of metalloprotease cleavage sites on membrane proteins, we employed a quantitative terminomics workflow consisting of shedding activation with phorbol 12-myristate 13-acetate (PMA) (Huovila et al., 2005), broad-spectrum metalloprotease inhibitor BB-94 (batimastat) treatment, SCX-based terminal peptide enrichment (Alpert et al., 2010; Chang et al., 2020; Helbig et al., 2010) (Figure S1A-D), TMT labeling and nanoLC/MS/MS measurement (Figure A-C).

We investigated secreted protein fractions across a panel of ten human cell lines stimulated with PMA following BB-94 treatment (Figure 1A, B). Samples were prepared in triplicate for each condition. The digest of 10 μg protein per replicate was subjected to terminal peptide enrichment: thus, in total only 1.2 mg of protein was utilized in the whole of this study. Terminal peptide enrichment was performed by SCX-StageTip (Adachi et al., 2016; Rappsilber et al., 2007), as described in the legend of Figure S1C, D and in the Methods section. Note that N-terminal peptides were TMT-labeled after enrichment, since the TMT tag alters the charge of peptides and affects enrichment efficiency (Figure 1C). For C-terminomics, the flow-through and the 0.5% TFA-eluted fraction were separately collected and subjected to LC/MS/MS (Figure S1D). Triplicates of controls and of BB-94-treated samples were multiplexed using 6-plexed TMT reagents (Figure 1D). We analyzed the TMT-labeled terminal peptides by high-resolution Orbitrap mass spectrometry considering a wide search space in semi-specific search mode (Niedermaier and Huesgen, 2019). Semi-specific search was performed using the Andromeda search engine on MaxQuant (Cox and Mann, 2008; Cox et al., 2011) against the SwissProt human protein database, including isoform sequences. TMT-reporter intensities were normalized by the trimmed mean of M values (TMM) method (Robinson and Oshlack, 2010).

**Figure 1.**
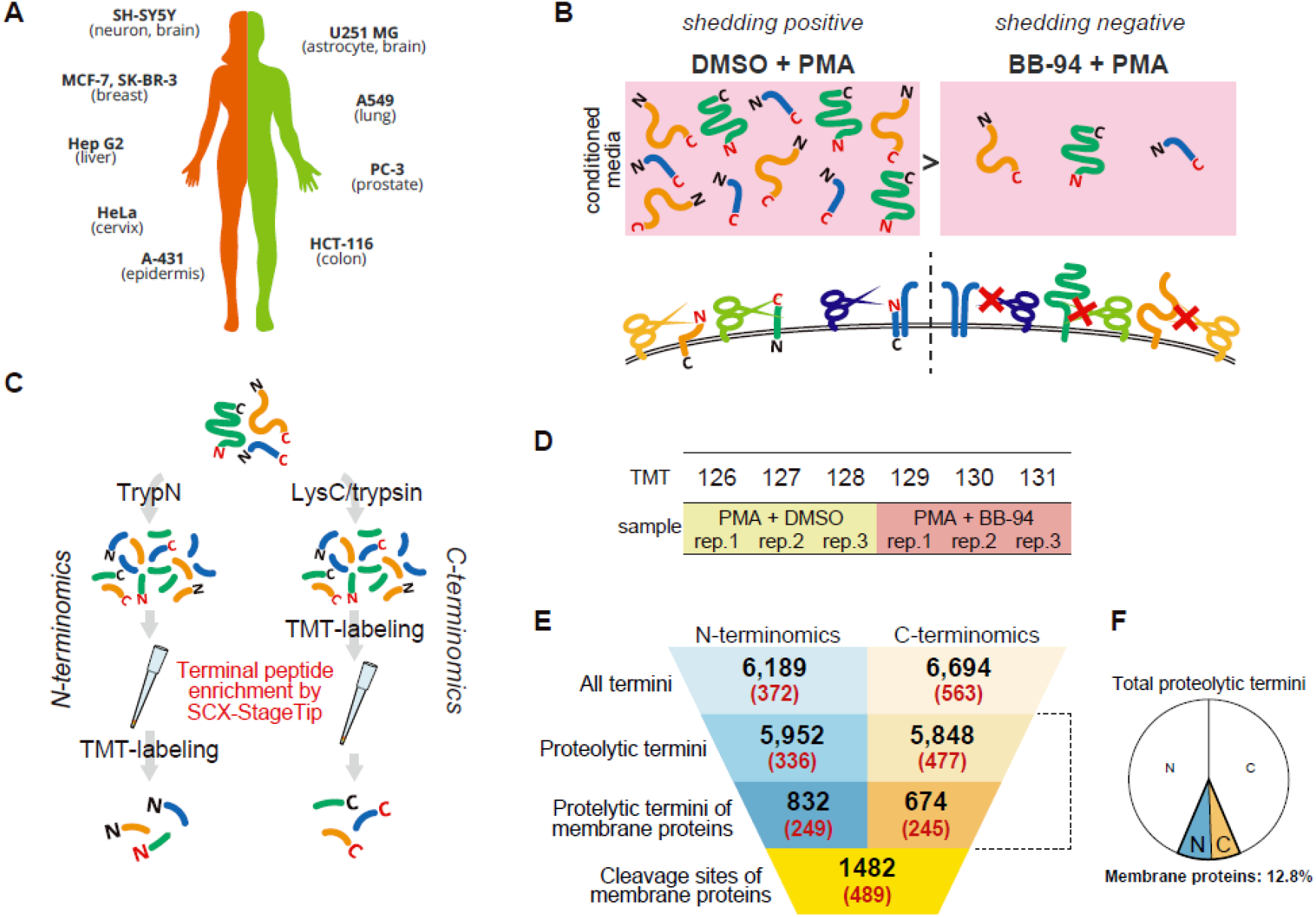
Experimental design and quantitative terminomics workflow for large-scale analysis of cleavage sites by metalloproteases. (A) List of investigated cultured cell lines (10 human cancer cell lines). (B) Experimental design. Cells were treated with DMSO or BB-94 for 1 h, followed by PMA treatment for 1 h. Triplicate samples were prepared for each condition. (C) Workflow of sample preparation. Proteins were digested with either TrypN or LysC and trypsin. Note that N-terminal peptides were TMT-labeled after terminal peptide enrichment, while C-terminal peptide enrichment was performed following TMT-labeling. See figure S1C, D for details. (D) Triplicate shedding-positive samples (PMA + DMSO) and shedding-negative samples (PMA + BB-94) were labeled as shown with TMT channels and form a TMT-6plex. (E) Summary of the number of identified termini and resulting cleavage sites. The parenthesized numbers show those significantly downregulated by BB-94. (F) Pie chart depicting the ratio of proteolytic termini mapped on membrane proteins in total proteolytic termini. The white areas show non-membrane proteins, and the areas highlighted with colors show membrane proteins. N, N-termini; C, C-termini.

Our strategy led to the identification of 6,189 N-termini and 6,694 C-termini in total. These included 5,952 proteolytic N-termini and 5,848 proteolytic C-termini, which are non-native protein termini and might have been generated by proteolysis (Figure 1E, Table S1-S2). In C-terminomics, C-terminal peptides accounted for 38% and 13% on average of the identified peptides in the flow-through and the 0.5% TFA-eluted fractions, respectively (Figure S1E). Of identified proteolytic termini, 832 N- and 674 C-termini were derived from membrane proteins (14.0% and 11.5% of the respective proteolytic terminome), which were defined using the UniProtKB Keywords ‘transmembrane’ or ‘GPI-anchor’. Figure S2-3 summarizes the reproducibility of three replicates. For individual conditions, there was excellent reproducibility in the peak intensity of identified peptides, with Pearson correlation coefficients in the ranges of *R* >0.93 (0.98 on average) for N-terminomics and *R* >0.97 for C-terminomics (0.99 on average). We created volcano plots with truncation at the false discovery rate (FDR) of 0.05 and artificial within-groups variance (S_0_) of 0.1 (default parameters) using Perseus (Tyanova et al., 2016) (Figure S4, 5), which yielded 336 proteolytic N-termini and 477 proteolytic C-termini that were significantly downregulated upon BB-94 treatment (Figure 1E). Importantly, these included 249 proteolytic N-termini (74.1% of downregulated proteolytic N-termini) and 245 proteolytic C-termini (51.4% of downregulated proteolytic C-termini) mapped on membrane proteins, affording a total of 489 cleavage sites (Table S3). Note that some cleavage sites were presented by both proteolytic N- and C-terminal peptides. As our results show (Figure 1F), proteolytic terminal peptides derived from membrane proteins account for a small fraction of the total proteolytic terminal peptides, which is one reason why they are so difficult to detect with MS. Our workflow enabled efficient identification and quantification of both proteolytic N- and C-termini of membrane proteins, highlighting a major strength of our methodology for achieving deep analysis of membrane protein cleavage sites with a limited amount of material.

### BB-94 selectively downregulates proteolytic termini of membrane proteins in the secretome

The number of downregulated termini varied among investigated cell lines (Figure 2A). U-251 MG cells have the highest number of downregulated termini, while MCF-7 have the lowest number. First, we examined the subcellular distribution of downregulated proteins using Gene Ontology (GO) term enrichment analysis by DAVID (v6.8; https://david.ncifcrf.gov/) (Huang et al., 2009). We found strong enrichment for terms that evoke membrane proteins, such as plasma membrane and cell surface (Figure 2B). In addition, in most samples, except the N-terminome of SK-BR-3, proteolytic termini of membrane proteins were downregulated by BB-94, while the abundances of proteolytic termini of non-membrane proteins were not changed (Figure 2C, D). These results indicate that BB-94 broadly and selectively targets the cleavage of membrane proteins, which would be useful for studying metalloprotease-dependent shedding. In addition, the provided cleavage site information in living cells should facilitate an understanding of the mechanism of action of BB-94.

**Figure 2.**
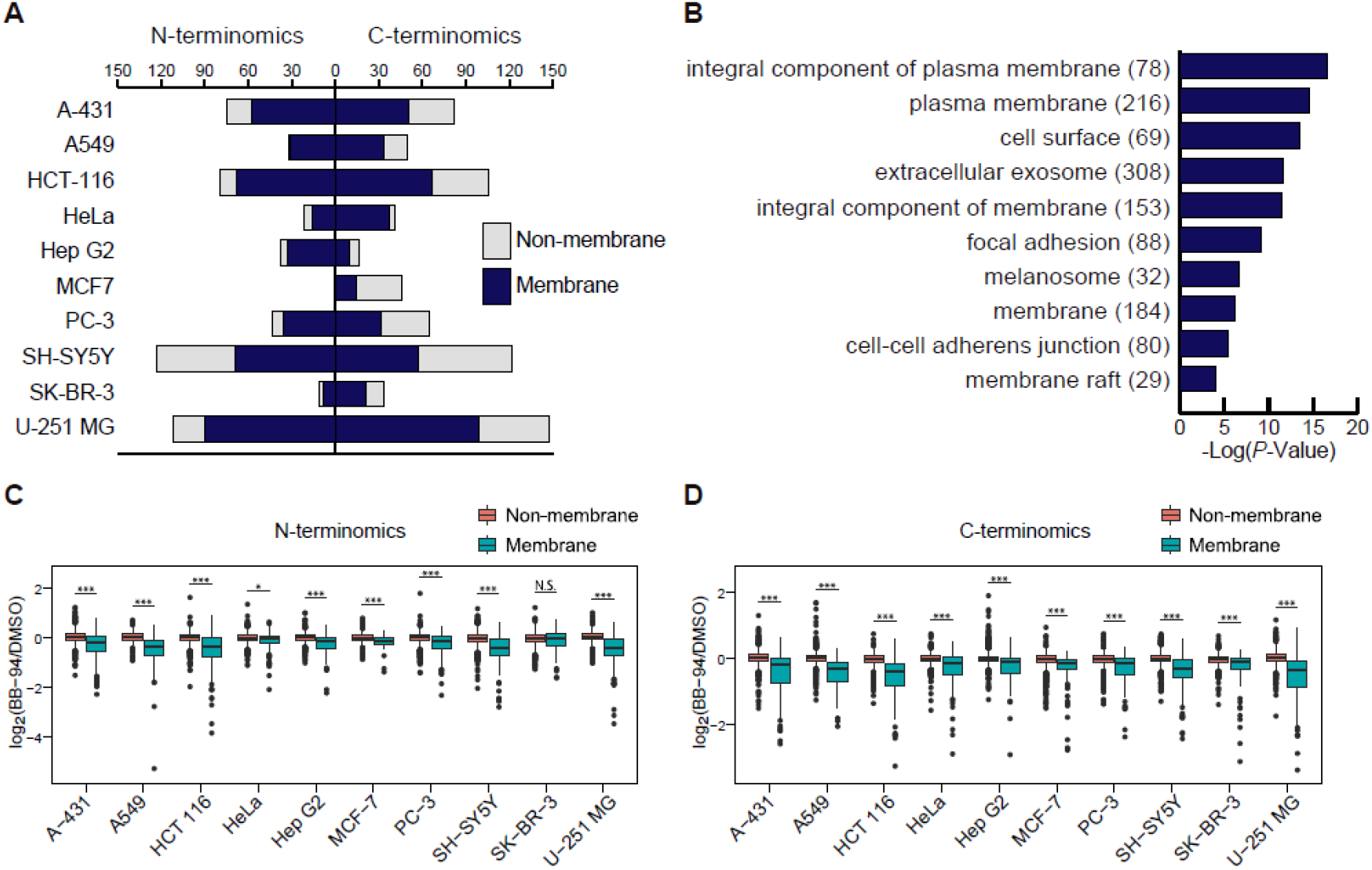
Overview of the BB-94-downregulated proteolytic terminome in the supernatant. (A) Distribution of the number of downregulated proteolytic termini mapped on membrane proteins and non-membrane proteins in the respective cell lines. (B) Enrichment analysis on GO term cellular components performed by DAVID (v8.0). The top ten significant terms are shown, with Bonferroni-adjusted *P*-values. The parenthesized numbers show the number of proteins. (C, D) The log_2_-transformed ratios (BB-94/DMSO) of proteolytic termini are compared between membrane proteins and non-membrane proteins in the respective cell lines. The *P*-values were calculated with the Wilcoxon rank sum test and a Bonferroni adjustment was applied. ***, *P*<0.005; **, *P*<0.001; *, *P*<0.05; N.S., not significant.

### Topological analysis of downregulated membrane proteins and positional analysis of the cleavage sites

We found that some of the downregulated membrane protein cleavage sites are localized at or around signal peptide cleavage sites (Figure S6A). We excluded 56 sites within ±5 amino acids from the signal peptide cleavage sites from further analyses in this study. Notably, the sequences around the signal peptide cleavage sites showed overrepresention of AxA at P3-P1 (three to one amino acids upstream of the cleavage sites), which is the motif of canonical signal peptidases (Figure S6B) (Paetzel et al., 2002). Topological analysis of the remaining 433 sites revealed that single-pass type I membrane proteins, which have their N-termini in the extracellular region, are the majority (378 sites, 87.3%), followed by GPI-anchor proteins (21 sites, 4.8%), singlepass type II membrane proteins, which have the C-termini in the extracellular region (17 sites, 3.9%), and multi-pass membrane proteins, which span the membrane more than once (10 sites, 2.3%) (Figure 3A). We identified five BB-94-downregulated cleavage sites on the extracellular domain of three multi-pass membrane proteins (cadherin EGF LAG seven-pass G-type receptor 2, G-protein coupled receptor 126, and Immunoglobulin superfamily member 1), although how these proteins were secreted into the culture media is unclear.

**Figure 3.**
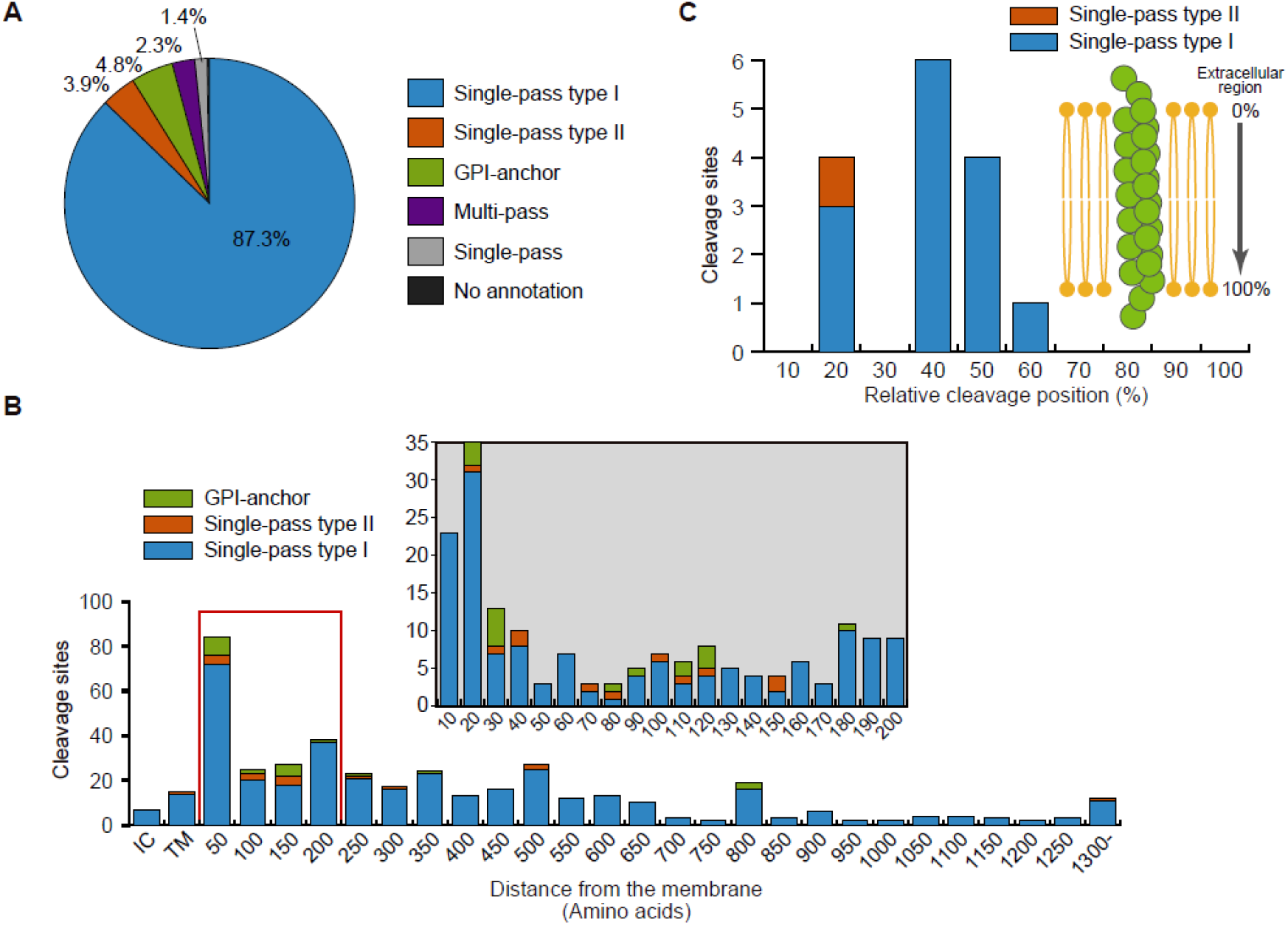
Topological analysis of downregulated membrane proteins and positional analysis of the cleavage sites. (A) Percentages of the various topologies of membrane proteins identified with downregulated cleavage sites. (B) The distribution of the distance (number of amino acids) from the transmembrane domain or GPI-anchor site to the cleavage site is depicted for single-pass type-I and type-II membrane proteins and GPI-anchored proteins. The section surrounded with a red box is shown in detail above. The cleavage sites mapped on the intracellular region are toward the minus direction. TM, cleavage sites are within the transmembrane domain. (C) Relative cleavage positions of the cleavage sites within the transmembrane domain of single-pass type I and type II membrane proteins. Values represent the length (number of amino acids) from the extracellular region to the cleavage site, divided by the total length (number of amino acids) of the transmembrane domain.

We examined the cleavage positions of single-pass type I and type II membrane proteins and GPI-anchor proteins. Of the 416 cleavage sites, 394 sites (94.7%) were distributed on the extracellular domain, confirming that BB-94 targets ‘ectodomain shedding’ (Figure 3B). Notably, we found that metalloprotease-regulated proteolysis occurred in close proximity to the cell surface: 174 sites (44.2%) were within 200 amino acids from the transmembrane domain or GPI-anchor site, which evidently suggests that large portions of the extracellular domains are released by shedding. This trend is consistent with previous findings on individual shedding substrates (Hinkle et al., 2004; Shirakabe et al., 2011; Zheng et al., 2004), and thus validates our results. On the other hand, 15 BB-94-downregulated sites were mapped within the transmembrane domain (Figure 3B), including three APP cleavage sites considered to be cleaved by γ-secretase (corresponding to amyloid β 37, 38, and 40) (Takami et al., 2009) (Figure S6C), and this suggests that our dataset includes cleavage sites generated downstream from metalloprotease-dependent shedding (Brown et al., 2000; Edwards et al., 2008; Reiss and Saftig, 2009). Interestingly, these downregulated cleavage sites inside the transmembrane domain were likely to be located at the extracellular side (Figure 3C). γ-Secretase cleaves APP in a stepwise manner, in which APP is first cleaved at the membrane-cytoplasm boundary, followed by successive tri- or tetrapeptide trimming (Takami et al., 2009). Consequently, the resulting terminus may be positioned at the extracellular side. Notably, in addition to APP, we identified five downregulated proteins (ALCAM in A431 and HCT 116; GLG1 in HCT 116; PIGR in A431; SDC1 in A431, A-549, HCT 116, HeLa, PC-3, SH-SY5Y, SK-BR-3, and U-251 MG; SDC4 in HCT 116) with cleavage sites on the extracellular domain and inside the transmembrane domain in the same cell line (Table S3). These proteins could have roles in bi-directional signal transduction across the cell membrane (Beard et al., 2019; Reiss and Saftig, 2009; Weber and Saftig, 2012).

### Functional characterization of ectodomain shedding

We functionally characterized the downregulated cleavage sites on the extracellular domain as metalloprotease-regulated shedding cleavage sites (394 cleavage sites on 119 proteins; Figure 3B). DAVID enrichment analysis (Huang et al., 2009) for GO term biological processes revealed significant enrichment for cell adhesion proteins (Figure 4), which is consistent with a previous study (Tsumagari et al., 2017). In addition, we found enrichment of proteins related to cell migration, including CD44, whose shedding is important for CD44-dependent cell migration (Nagano and Saya, 2004). Analysis of GO terms of molecular function showed that shedding targets central components of signal transduction, such as receptors and protein tyrosine kinases/phosphatases. Importantly, we observed that shedding liberates a large portion of the extracellular domain (Figure 3B), which is the main functional region of membrane proteins, on a proteomic scale. Thus, overall, our results indicate that shedding regulates fundamental biological events by modulating membrane protein functions.

**Figure 4.**
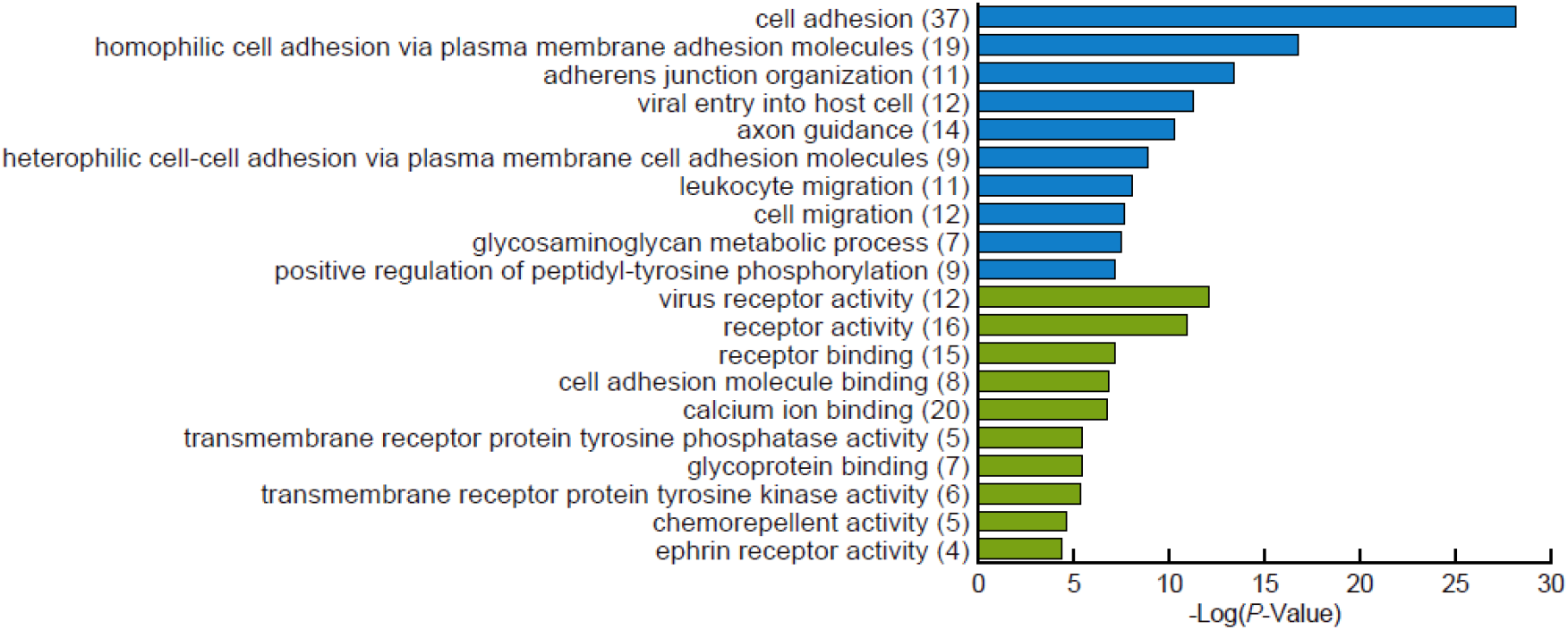
Functional analysis of metalloprotease-regulated shedding substrates. DAVID enrichment analysis of metalloprotease-regulated shedding substrates according to GO term biological processes (blue) and molecular functions (green). The top ten significant terms are shown, with Bonferroni-adjusted *P*-values. The parenthesized numbers show the numbers of proteins.

### Cell-type-specificity of shedding

Differently processed proteoforms can exhibit distinct physiological properties (Chow et al., 2010; Hu et al., 2006; La Marca et al., 2011; Willem et al., 2006). Whereas most BB-94-downregulated membrane proteins contain one cleavage site, interestingly no less than 65 cleavage sites were identified on neuronal cell adhesion molecule (NrCAM) (Figure 5A). This indicates that proteolysis can produce an impressive number of proteoforms, and may critically contribute to the complexity of the human proteome.

**Figure 5.**
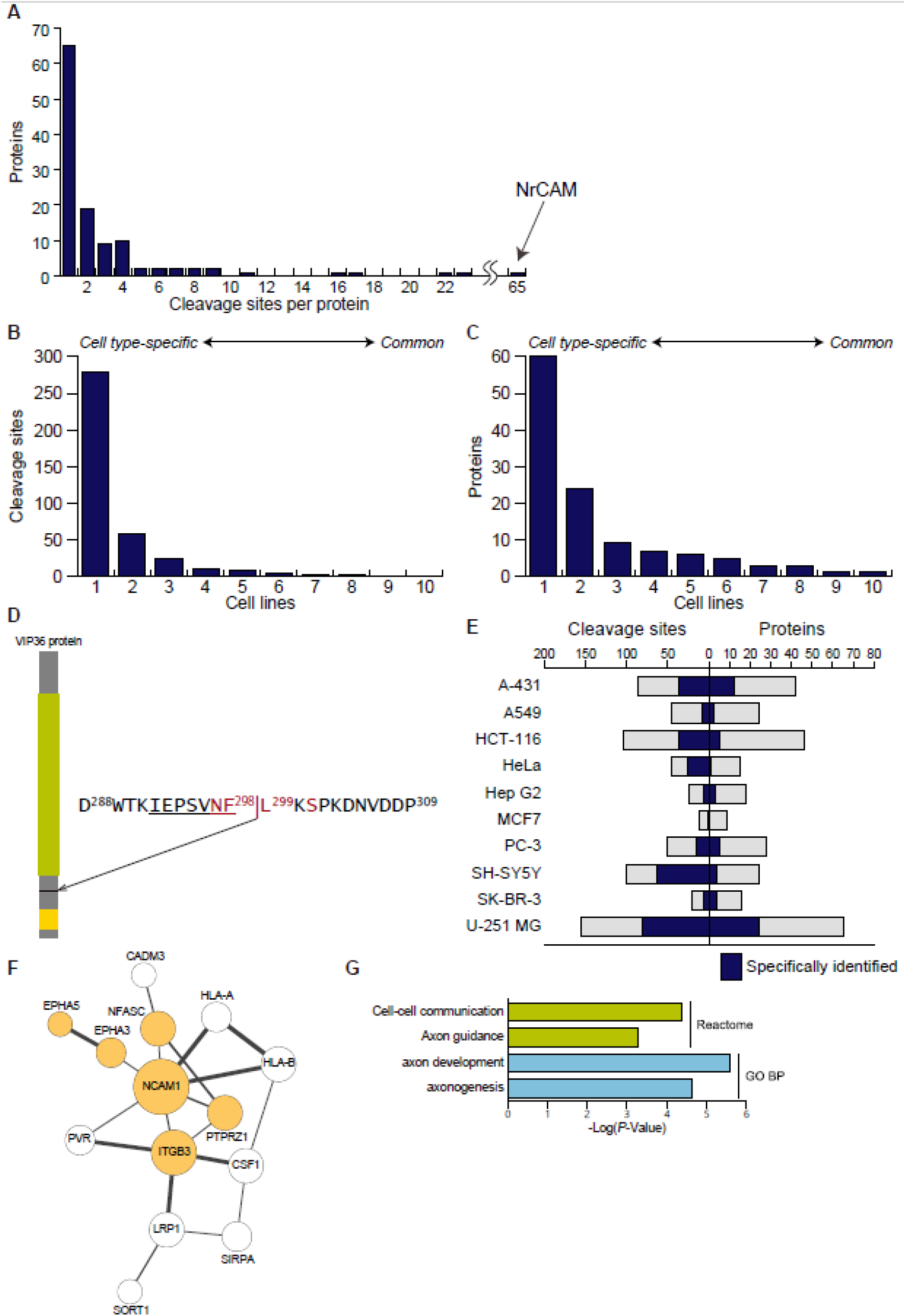
Occurrence and cell type-specificity of shedding. (A) The distribution of the number of cleavage sites per protein is depicted for single-pass type-I and type-II transmembrane proteins and GPI-anchored proteins. (B, C) Histograms depicting the number of shedding substrate identified in 1 to 10 cell lines at the cleavage site level (B) and at the protein level (C). (D) VIP36 cleavage site. The amino acids that were previously suggested to be essential for cleavage are highlighted in red (Shirakabe et al., 2011). The sequence of the peptide identified in this study is underlined. (E) Distribution of the number of metalloprotease-regulated cleavage sites in the respective cell lines. The number of cleavage sites specifically identified in each cell type is highlighted in color. (F, G) Protein interaction network of shedding substrate proteins specifically identified in U-251 MG cells (F). Proteins without any interactions are excluded. Significantly enriched reactome and GO terms of interest are shown with the Benjamini-adjusted *P*-values (G), and the proteins annotated with these terms are highlighted in color (F). All analyses were performed by STRING (v11). The network was visualized using Cytoscape (v3.8.0). Node size reflects the number of connections (direct edges), and edge line width reflects the combined score calculated by STRING.

A large population of the identified shedding substrates was cell-type-specific rather than globally identified, at both the cleavage site and protein levels (Figure 5B, C), suggesting that shedding occurs depending on the individual functions of the cells. Sixty proteins were found in one cell line, while only one protein, 36-kDa vesicular integral-membrane protein VIP36 coded by the LMAN2 gene, was found commonly in all investigated cell lines. VIP36 is a validated shedding substrate (Shirakabe et al., 2011). Intriguingly, the VIP36 protein was cleaved at the same site (F298-L299) across all analyzed cell lines. In the previous study, western blotting in combination with amino acid substitution at expected cleavage sites failed to identify the precise cleavage site of VIP36 (Shirakabe et al., 2011), but the site we identified here is positioned just within the expected region (Figure 5D), which again validates our result and underscores the strength of our strategy. VIP36 is a lectin domain-containing transmembrane protein that functions as a cargo receptor transporting glycoproteins. While shedding of the VIP36 protein plays a critical role in phagocytosis in Raw 264.7 cells (Shirakabe et al., 2011), the role of VIP36 shedding in other cell types is unknown.

We identified cell-line-specific shedding substrate proteins in nine cell lines, except for MCF-7. Astrocytoma-derived U-251 MG cells yielded the highest number of substrates (Figure 5E), and 24 proteins were uniquely identified in U-251 MG cells. To investigate the functions of these U-251 MG cell-specific substrates, we investigated protein-protein interactions of these substrate proteins using the STRING database (v11.0; https://string-db.org/) (Szklarczyk et al., 2019), which resulted in significantly more interactions than would be expected for a random protein set of similar size with a *P*-value of 6.13e-11 (calculated by STRING) (Figure 5F). Interestingly, enrichment analysis revealed that these U-251 MG cell-specific substrates are related to typical functions of neurons, such as axon development and axon guidance, rather than those of astrocytes (Figure 5G). This may indicate that shedding contributes to cellular communication across different cell types in the central nervous system.

### Evaluation of metalloprotease-regulated shedding cleavage sites using PWM scoring

As mentioned above, our results include cleavage sites downstream of the initial metalloprotease cleavages. Therefore, to identify the direct cleavage sites by metalloproteases with greater confidence, we evaluated the identified ectodomain cleavage sites of type-I membrane proteins and GPI-anchored proteins, whose N-terminal regions are located on the extracellular surface, using position weight matrix (PWM) scoring (Imamura et al., 2017) with the accumulated substrate information in the MEROPS protease database (v.12.1, https://www.ebi.ac.uk/merops/index.shtml) (Rawlings et al., 2018). Although ADAM17 and ADAM10 are major sheddases (Huovila et al., 2005), the numbers of their registered substrate cleavage sites are much fewer than those of other metalloproteases due to the relative absence of studies on their cleavage sites, despite their physiological importance. We concatenated *in vitro* substrate information to the MEROPS-registered substrates (Tucher et al., 2014), which yielded 225 and 381 cleavage site sequences for ADAM10 and ADAM17, respectively. In addition we further selected 14 sheddases (matrix metalloprotease (MMP) −2, −3, −7, −8, −9, −12, −13, MT1-MMP, legumain, meprin β, cathepsin S, cathepsin L, furin and proprotein convertase subtilisin/kexin 7 (PCSK7)) from various families, based on following criteria: described as canonical or part-time sheddases in a recent review (Lichtenthaler et al., 2018); having a number of registered substrates in MEROPS greater than one hundred.

We computed substrate sequence PWMs for these individual sheddases (Figure 6A, S7). Importantly, individual sheddases showed distinct substrate preference signatures. Overall, most metalloproteases, except meprin β, commonly exhibit a preference for hydrophobic residues such as Leu and Ile at the P1’ position (one amino acid downstream of the cleavage site), and for Pro and other hydrophobic residues at the P3 position (three amino acids upstream of the cleavage sites); these trends are consistent with previous reports (Eckhard et al., 2016; Tucher et al., 2014). ADAM10 characteristically exhibits a preference for bulky hydrophobic residues such as Tyr, Trp and Phe, which reflects their structural differences (Seegar et al., 2017). Furin and PCSK7 showed known RxxR or RxKR cleavage motifs at P4-P1 (Seidah et al., 2013). Legumain and meprin β showed preferences for Asp/Asn at P1 and acidic residues at PT, respectively. Given that the PWMs are validated, they can provide a basis for scoring identified putative shedding sites of the respective sheddases.

**Figure 6.**
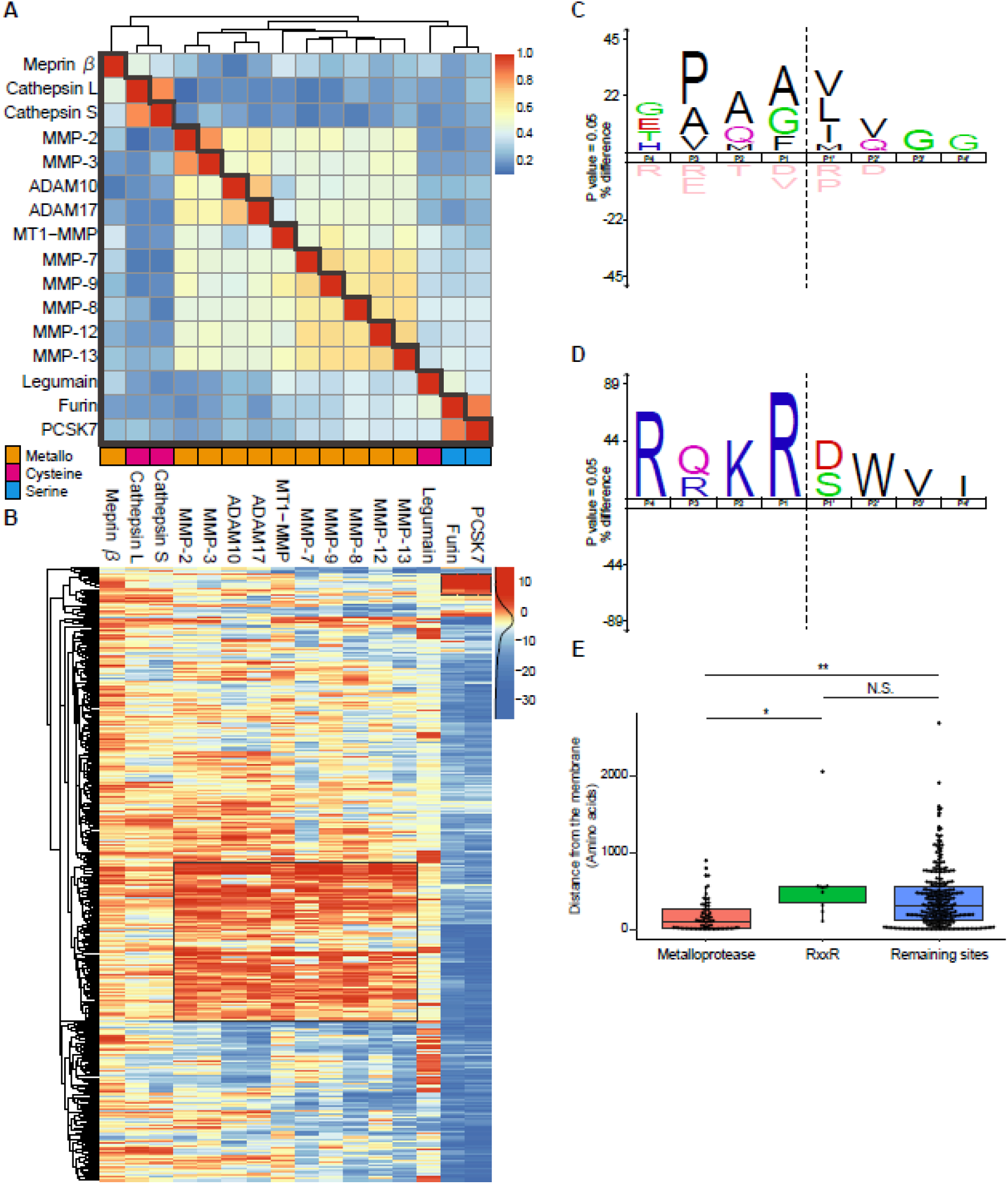
PWM scoring of downregulated cleavage sites. (A) Heatmap depicting the cosine similarity between PWMs for 16 selected sheddases shown in figure S7. High similarity is highlighted in red, while low similarity is highlighted in blue. Sheddases are ordered by clustering based on the cosine distances (1 – cosine similarity). (B) Hierarchical clustering analysis of PWM scores. High-scored sites are highlighted in red, while low-scored sites are highlighted in blue. A cluster of cleavage sites high-scored for metalloproteases (metalloprotease cluster) and a cluster for furin and PCSK7 (RxxR cluster) are enclosed with solid-line and dashed-line boxes, respectively. (C, D) Sequence logos of cleavage sites in the metalloprotease cluster (C) and in the RxxR cluster (D) generated by iceLogo (https://iomics.ugent.be/icelogoserver/). Dashed line shows the cleavage site. (E) The distances from the transmembrane domain or GPI-anchor site to the cleavage site are compared between the metalloprotease cluster, the RxxR motif cluster, and the remaining sites. The *P*-values were calculated with the Wilcoxon rank sum test and Bonferroni adjustment was applied. **, *P* = 1.26e-07; *, *P* = 8.82e-4; N.S., not significant.

We calculated PWM scores between more than 6000 relationships (16 sheddases x 378 cleavage sites) (Figure 6B, Table S4). Clustering analysis revealed a group of 86 cleavage sites high-scored for metalloproteases, including the α cleavage site of APP mediated by ADAM10 (Figure 6B, C; Table 1). This cluster also included the VIP36 cleavage site that was commonly identified in the analyzed cultured cells (Figure 5D). Notably, although these proteins were previously known to undergo shedding, in many cases, the exact cleavage sites that we identified here have not been reported. We also found another cluster, which involving the RxxR motif of furin and PCSK7 (Figure 6B, D). These serine proteases are considered as part-time sheddases, which mainly function as proprotein convertases, but can additionally participate in shedding (Lichtenthaler et al., 2018). The downregulation of these cleavages upon BB-94 treatment might be a secondary or later event following the inhibition of metalloproteases, rather than being due to direct inhibition by BB-94.

**Table 1.**
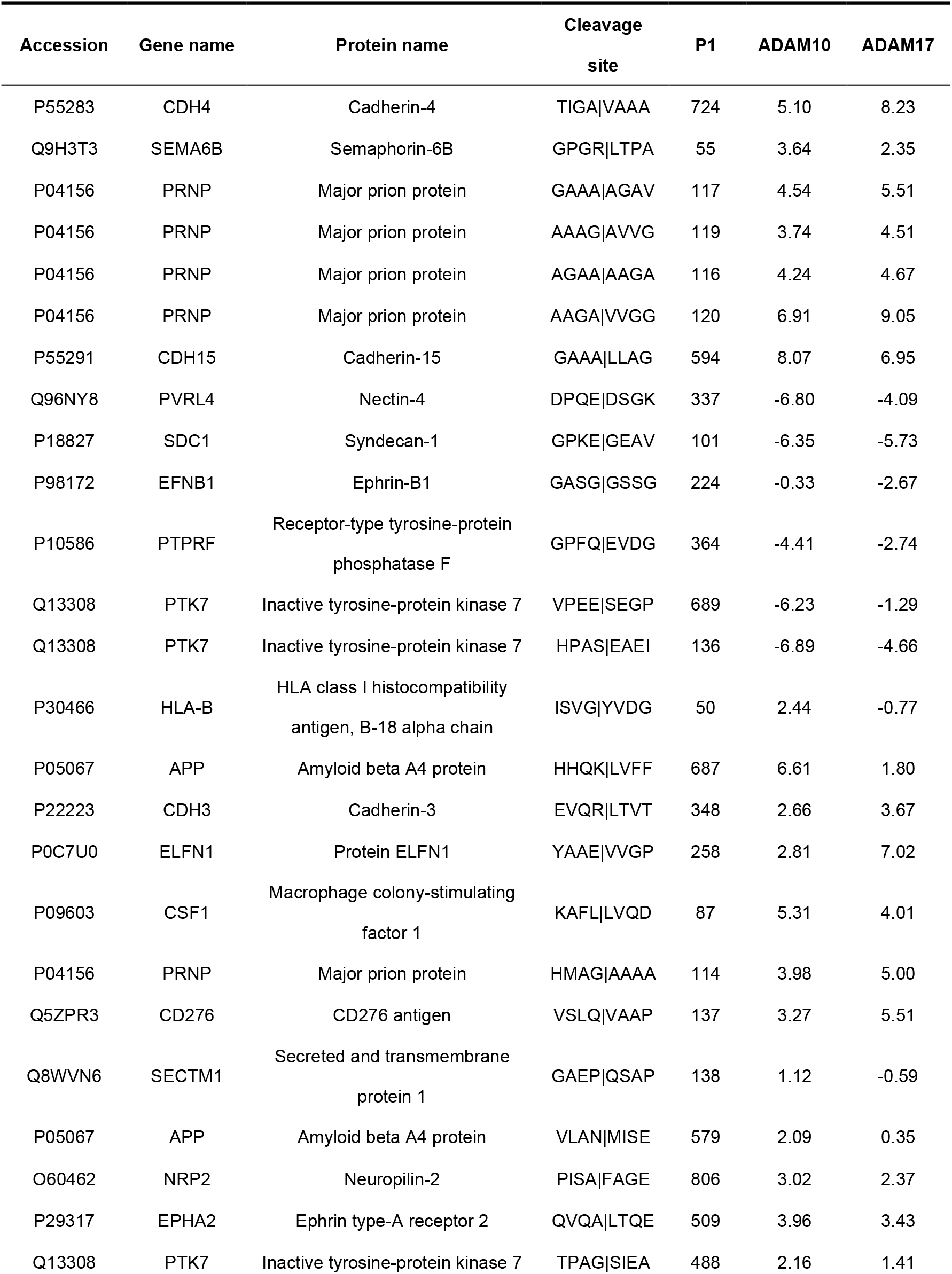
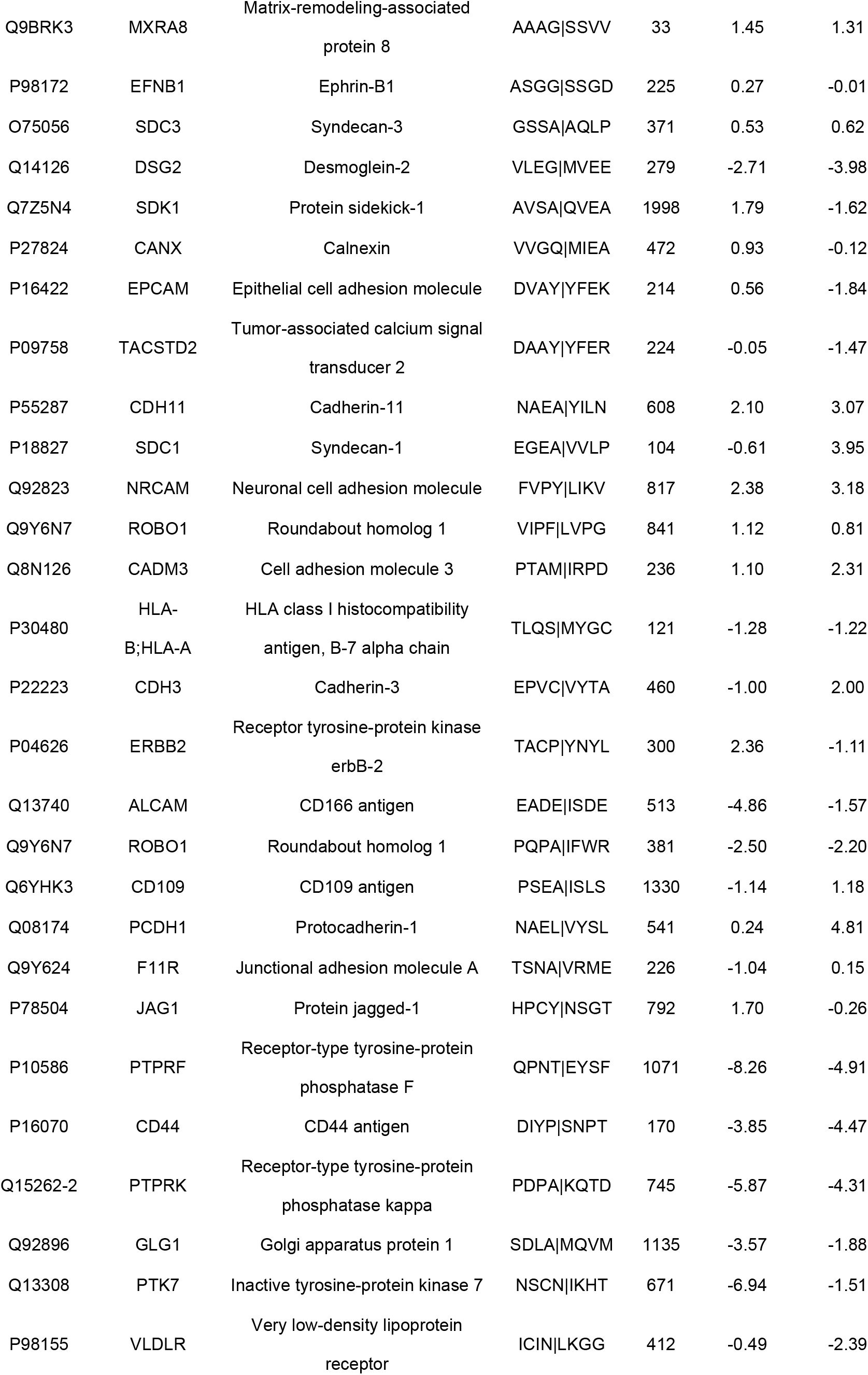
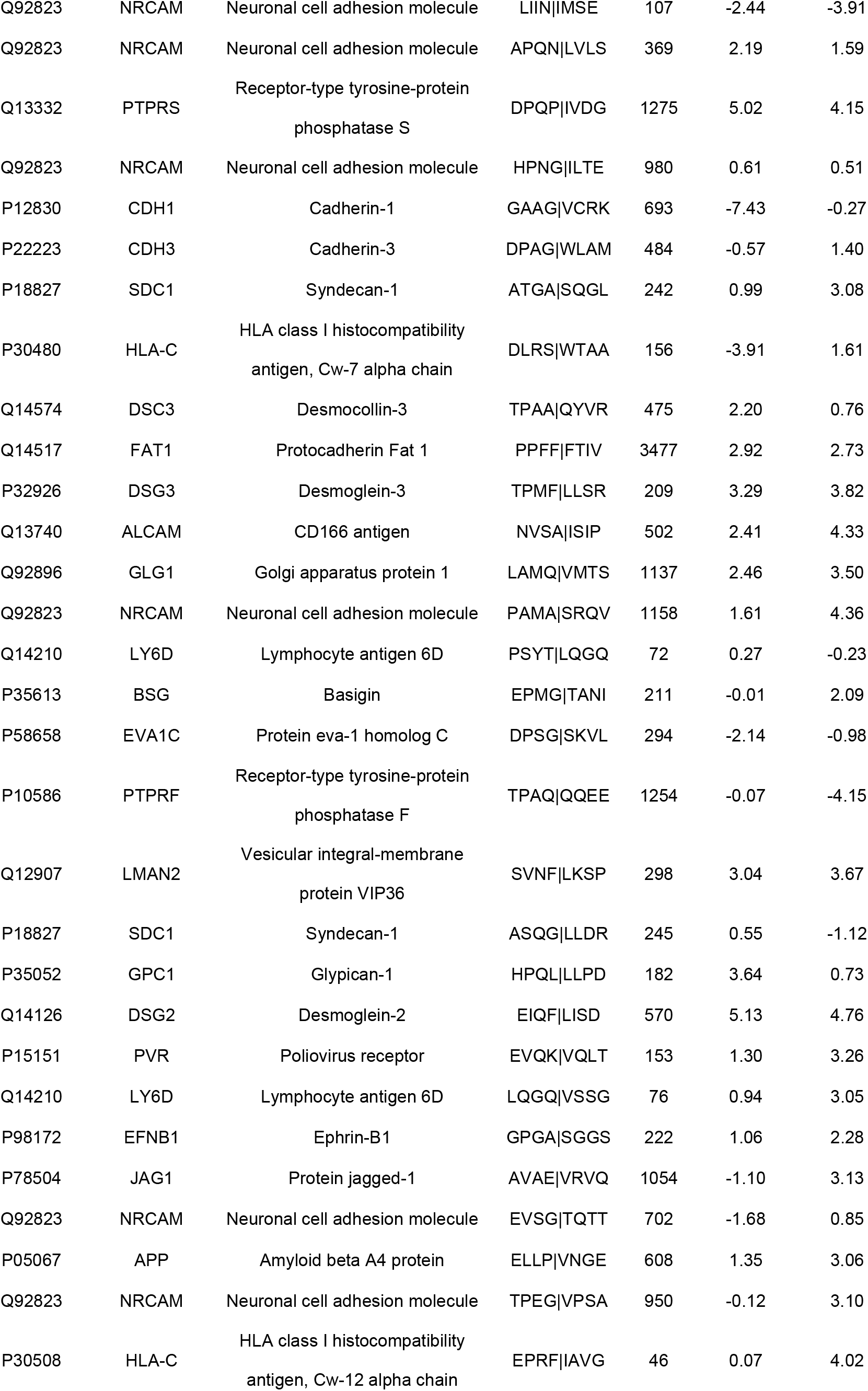
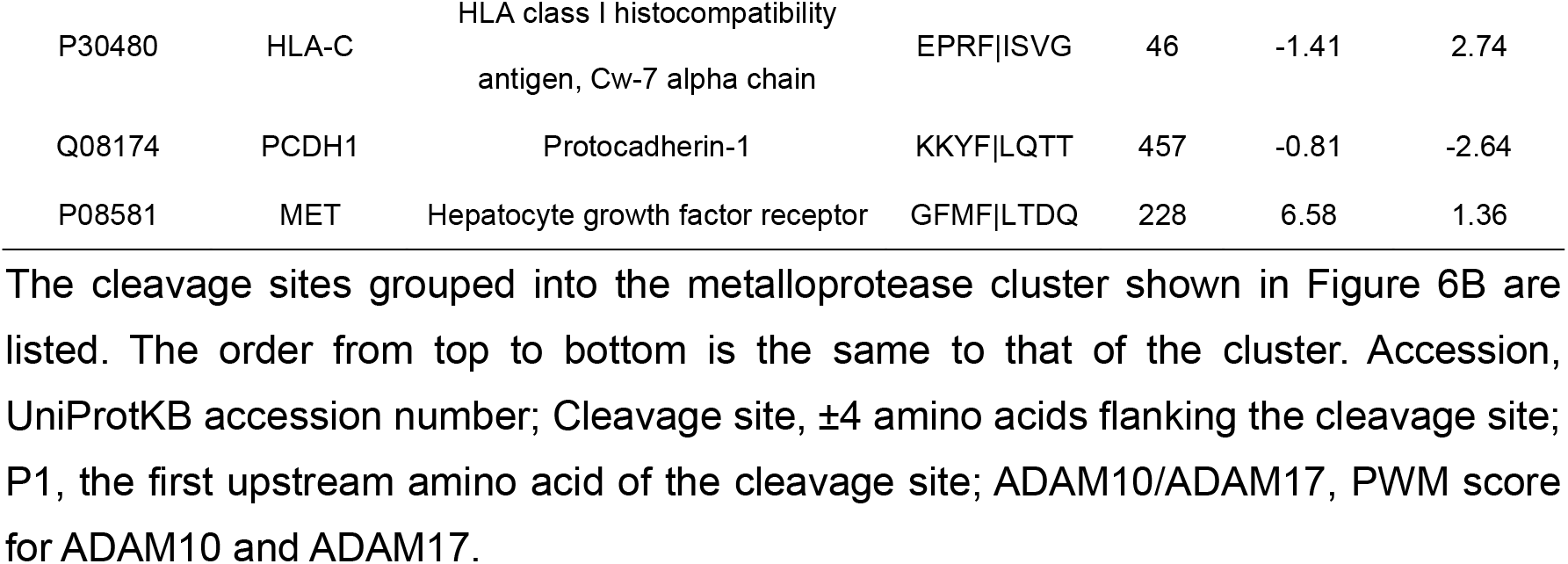
List of cleavage sites grouped into the metalloprotease cluster.

Comparison of the distances from the transmembrane domain to the cleavage site in the metalloprotease cluster, the RxxR cluster, and the remaining sites revealed that the cleavage sites in the metalloprotease cluster are located closer to the cell surface (Figure 6E). Furthermore, the cleavage sites close to the membrane (<200 amino acids in Figure 3B) are significantly enriched in metalloprotease cluster cleavage sites with the *P*-value of 2.5e-3 (Fisher’s exact test). These results confirm that metalloproteases shed their substrates at positions close to the membrane.

## Discussion

Here, we have demonstrated that SCX-based N- and C-terminal peptide enrichment in combination with high-resolution mass spectrometry allows reproducible quantification of thousands of protein termini derived from membrane proteins in culture supernatant, with limited amounts of materials. As our protocol of C-terminomics consists of simple steps, including isobaric tagging and tip-based SCX, it should prove to be a practically useful platform not only for identification of shedding substrates in the supernatant, but also for conventional C-terminomics of cells and tissues. The results were validated by the observation of membrane-protein-selective downregulation by BB-94. Broadly speaking, shedding occurs in the vicinity of the cell surface, thus liberating a large portion of the extracellular domain. Also, shedding targets the main components of signal transduction including protein tyrosine kinases/phosphatases and receptors. We successfully identified the cleavage site of VIP36, which could not be fully clarified by amino acid substitution and western blotting studies (Shirakabe et al., 2011), and interestingly this was the only cleavage site commonly identified across the 10 investigated cell lines. The importance of investigating diverse cell types is highlighted by the substantial differences of shedding pattern across the analyzed cell lines. We focused on the function of U-251 MG cell-unique shedding and found that the released proteins are important for the environment, rather than the cells themselves. Overall, our results provide further support for the idea that shedding contributes to cellular communication. Many of the identified putative substrate proteins were previously reported to undergo shedding, and here we have unveiled the exact cleavage sites. We wish to emphasize that this is the first study in which native shedding is overviewed at the cleavage site level on a proteome-wide scale across multiple cell types.

Our data set contains cleavage sites generated downstream of metalloprotease-dependent shedding. In addition, all fragments may be further processed by a wide range of extracellular endo- and exopeptidases. We therefore evaluated identified metalloprotease-regulated cleavage sites by PWM scoring, which we previously employed to evaluate protein kinase-phosphorylation site relationships (Imamura et al., 2017). We successfully found cleavage sites high-scored for metalloproteases, which could represent the direct substrates of the metalloproteases. In addition, we found a cluster high-scored for furin and PCSK7, which have a clear cleavage motif, RxxR. Proteolytic processing by these proteases is mainly involved in the maturation of membrane proteins. These proteases might be activated as a result of drastic upregulation of membrane protein turnover by PMA-activated shedding. We observed distinct signatures of PWMs even for ADAMs and MMPs, which have extremely similar preferences. In the future, accumulating knowledge of cleavage sites should enable even more powerful approaches, such as machine learning, for confident prediction of sheddase-cleavage site relationships.

Our data set provides a useful resource to support hypothesis-driven studies on shedding. It seems clear that a technological breakthrough in proteomic identification of shedding cleavage sites is needed, not only from a scientific viewpoint, but also for therapeutic purposes, given the involvement of shedding in various diseases, including cancer, cardiovascular disease, inflammation, and neurodegeneration (Edwards et al., 2008; Herrlich and Herrlich, 2017; Weber and Saftig, 2012). It has been suggested that cells shed different membrane proteins depending upon external stimulation (Dang et al., 2013; Huovila et al., 2005), i.e., cells selectively shed substrates in response to different stimuli. However, how shedding is made specific has long been enigmatic. Most shedding is mediated by ADAM10 and ADAM17, and these sheddases share many substrates (Gooz, 2010; Huovila et al., 2005). In addition, sheddases generally have broad amino acid preferences, and do not require a specific amino acid at any position surrounding the cleavage site. Consequently, it is extremely difficult to understand the basal mechanisms that determine shedding targets. Our strategy presented here enables systematic investigation of changes of shedding at the site level and should facilitate work to identify critical factors that regulate shedding by means of proteomics studies.

## Acknowledgments

We thank Eito Yamamoto (Kyoto University) for assisting in the development of the C-terminal peptide enrichment protocol, and Atsuko Sehara-Fujisawa (Kyoto University) for providing SH-SY5Y cells. K.T. was supported by a fellowship for young scientists from the Japan Society for the Promotion of Science (JSPS). This work was supported by the JST Strategic Basic Research Program, CREST (grant No. 18070870), and by a JSPS Grant-in-Aid for Scientific Research (No. 17H05667).

## Author Contributions

Conceptualization, K.T. and Y.I.; Methodology, K.T. and C-H.C.; Formal analysis, K.T.; Investigation, K.T.; Resources, K.T.; Writing – Original Draft, K.T.; Writing – Review & Editing, Y.I.; Visualization, K.T.; Supervision, Y.I.; Project Administration, Y.I.; Funding Acquisition, K.T. and Y.I.

## Methods

### Cell culture

Basically, cells were cultured in Dulbecco’s modified Eagle’s medium (DMEM; FUJIFILM Wako) supplemented with 10% fetal bovine serum (Gibco), 100 U/mL penicillin and 100 μg/mL streptomycin (FUJIFILM Wako) on 15 cm culture dish at 37°C in a humidified incubator with 5% CO_2_. PC-3 cells were cultured in Roswell Park Memorial Institute 1640 medium (RPMI1640; FUJIFILM Wako) with the same supplements as listed above. SH-SY5Y cells and U-251 MG cells were cultured in DMEM/Ham’s F-12 (FUJIFILM Wako) including non-essential amino acids (FUJIFILM Wako) and the same supplements as listed above. We did not perform specific authentication of the cell lines used in this study.

Nine dishes of 90% confluent cells per condition were prepared. Supernatants of three dishes were pooled during supernatant collection to make a biological replicate.

### Inhibitor treatment and supernatant preparation

Cells were washed with phosphate-buffered saline (PBS; FUJIFILM Wako) three times and incubated for 1 h with Hanks’ balanced salt solution containing Ca^2+^ and Mg^2+^ (HBSS (+); FUJIFILM Wako) together with 10 μM BB-94 (Selleck) or DMSO (FUJIFILM Wako) for inhibitor pretreatment. Then, the buffer was replaced with fresh HBSS (+) containing 1 μM PMA (FUJIFILM Wako) and 20 μM BB-94 or DMSO, and the cells were incubated for 1 h. The supernatants were quickly collected on ice, and centrifuged at 7,180 *g* for 30 min to remove cell debris. Then 2 mM ethylenediaminetetraacetic acid (EDTA; DOJINDO), 2 mM ethyleneglycol bis(2-aminoethyl ether)-*N,N,N,N* tetraacetic acid (EGTA; DOJINDO), 1 mM phenylmethylsulfonyl fluoride (PMSF; FUJIFILM Wako) and 0.1x protease inhibitor cocktail (Sigma-Aldrich) were added to the supernatant to inhibit further proteolytic degradation, and the solution was stored at −80°C until use.

### Protein purification and digestion

Culture media were concentrated up to ~50-fold using an Amicon^®^ Ultra filter (3,000 NMWL; Merck), acidified with 2.5% trifluoroacetic acid (TFA; final concentration), and further concentrated using a SpeedVac (Thermo Fisher Scientific). Proteins were reconstituted with 100 μL water and purified by methanol-chloroform precipitation, for which the protein solution was successively mixed with 300 μL methanol, 75 μL chloroform, and 225 μL water and centrifuged at 21,000 *g* for 5 min. The resulting pellet was washed with 225 μL methanol and dissolved in phase-transfer surfactant (PTS) buffer (Masuda et al., 2009) containing 100 mM 2-amino-2-(hydroxymethyl)-1,3-propanediol hydrochloride (Tris-HCl, pH 8.5), 12 mM sodium deoxycholate (SDC; FUJIFILM Wako), 12 mM sodium *N*-lauroylsarcosinate (SLS; FUJIFILM Wako). The protein concentration was determined by means of bicinchoninic acid (BCA) assay (Thermo Fisher Scientific). For each condition, 10 μg of protein was reduced with 10 mM dithiothreitol (DTT; FUJIFILM Wako) and alkylated with 40 mM 2-chloroacetamide (CAA; Sigma-Aldrich). Digestion was performed according to the PTS protocol (Chang et al., 2020; Masuda et al., 2009). Briefly, for C-terminomics, the protein solution was 5-fold diluted with 50 mM ammonium bicarbonate, and proteins were sequentially digested with LysC (1:50, w/w; FUJIFILM Wako) for 3 h at 37 °C and trypsin (50:1, w/w; Promega) overnight at 37 °C. In the case of N-terminomics, following 10-fold dilution with 10 mM CaCl_2_, proteins were digested with TrypN (1:50, w/w; Protifi) overnight at 37 °C. Note that TrypN can be replaced with LysargiNase (Merck, Cat#EMS0008). Digestion was stopped with 0.5% TFA (final concentration), and the detergents were removed by liquid-liquid extraction using an equal volume of ethyl acetate. Peptides were purified with a reversed-phase SDB-XC StageTip (Rappsilber et al., 2007).

### TMT labeling

Peptides were dissolved in 5 μL of 200 mM 4-(2-hydroxyethyl)-1-piperazineethanesulfonic acid (HEPES; DOJINDO)-NaOH (pH 8.5), mixed with 0.1 mg of TMT reagents dissolved in 5 μL of acetonitrile (ACN), and agitated at room temperature for 1 h. Excessive reagents were quenched with 0.33% hydroxylamine (final concentration), and the solution was acidified with 1% TFA (final concentration). Note that N-terminal peptides were TMT-labeled after N-terminal peptide enrichment, and then 6-plexed peptides were mixed. ACN was evaporated using a SpeedVac. Labeled peptides were purified using a reversed-phase SDB-XC StageTip (Rappsilber et al., 2007).

### N-Terminal peptide enrichment

The TrypN digest of 10 μg protein was dissolved in 50 μL of N-loading buffer (2.5% formic acid (FA), 30% ACN) and loaded onto a StageTip with 16-gauge double-plug CationSR membranes (GL Sciences), which had been successively preconditioned with 50 μL methanol, 50 μL 80% ACN containing 0.1% TFA, 100 μL 500 mM ammonium acetate containing 30% ACN, and 300 μL N-loading buffer. Following peptide loading, the StageTip membranes were washed with 50 μL N-loading buffer, and the wash solution was collected together with the flow-through fraction as the N-terminal peptides-enriched fraction. This fraction was evaporated using a SpeedVac and TMT-labeled as described above.

### C-Terminal peptide enrichment

A mixture of 6-plexed TMT-labeled peptides was dissolved in 150 μL of C-loading buffer (0.15% TFA, 30% ACN), divided into three parts, and subjected to C-terminal peptide enrichment using three StageTips. A solution of peptides in 50 μL C-loading buffer was loaded onto a StageTips with 16-gauge double-plug CationSR membranes (GL Sciences), which had been successively preconditioned with 50 μL methanol, 50 μL 80% ACN containing 0.1% TFA, 100 μL 500 mM ammonium acetate containing 30% ACN, and 300 μL C-loading buffer. Subsequently, peptides were eluted with 0.5% TFA containing 30% ACN (0.5% TFA fraction). The flow-though fraction and the 0.5% TFA fraction were separately collected and evaporated.

### Liquid Chromatography/Tandem Mass Spectrometry

For N-terminal peptide analysis, one-third of the peptides was injected, and triplicate analyses were carried out (3 LC/MS/MS runs per cell line). For C-terminal peptide analysis, three sets of flow-through and 0.5% TFA fractions were subjected to single shot analysis (6 LC/MS/MS runs per cell line) (Figure S1C, D).

A nanoLC/MS/MS system comprising an UltiMate 3000RSLCnano pump (Thermo Fisher Scientific) and an Orbitrap Fusion Lumos tribrid mass spectrometer (Thermo Fisher Scientific) was employed. Peptides were injected by an HTC-PAL autosampler (CTC Analytics), loaded on a 15 cm fused-silica emitter self-packed with ReproSil-Pur C18-AQ (3 μm; Dr. Maisch), and separated by a linear gradient, that is, 5% B for 1 min, 5–15% B in 4 min, 15–45% B in 100 min, 40–99% B in 5 min, and 99% B for 10 min (Solvent A, 0.5% acetic acid; solvent B, 0.5% acetic acid in 80% ACN) at the flow rate of 500 nL/min. Peptides were ionized at 2,400 V. All MS1 spectra were acquired over the range of 375–1500 *m/z* in the Orbitrap analyzer (resolution = 120,000, maximum injection time = 50 ms, automatic gain control = 4e5). For the subsequent MS/MS analysis, precursor ions were selected and isolated in top-speed mode (cycle time = 3 sec, isolation window = 1.4 m/z), activated by higher-energy collisional dissociation (HCD; normalized collision energy = 38), and detected in the Orbitrap analyzer (resolution = 50,000, maximum injection time = 105 ms, automatic gain control = 1e5). Dynamic exclusion time was set to 30 sec.

### LC/MS/MS raw data processing

LC/MS/MS raw data were processed using MaxQuant (v.1.6.7.0) with the Andromeda search engine (Cox and Mann, 2008; Cox et al., 2011). Database search was implemented against the UniprotKB/SwissProt (2019_3) human database including isoform sequences concatenated with commonly observed contaminant protein sequences set in MaxQuant. Two analysis groups were made in MaxQuant, enabling one combined analysis for TrypN with N-terminal free semi-specificity and trypsin/P with C-terminal free semi-specificity. The following parameters were applied: 10-plexed TMT quantification at the MS2 level, precursor mass tolerance of 4.5 ppm, fragment ion mass tolerance of 20 ppm, and minimal peptide length of 7 amino acids. Cysteine carbamidomethylation was set as a fixed modification, while methionine oxidation and acetylation on the protein N-terminus were allowed as variable modifications. False discovery rates were estimated by searching against a reversed decoy database and filtered for <1% at the peptide-spectrum match, peptide and protein level.

### Protein inference

In terminomics, isoforms are often indistinguishable due to limited sequence coverage. Thus, we used the canonical isoform for bioinformatics analysis of membrane proteins, if the peptide can be derived from the canonical isoform. If not, we employed the leading protein given by MaxQuant.

### Peptide list processing and normalization of TMT-reporter intensities

Firstly, peptides derived from contaminant proteins such as keratins and bovine serum-derived proteins were excluded. We utilized peptides with no missed cleavages for analyses. TMT-reporter intensities were log_2_-converted and normalized in individual cell lines by the trimmed mean of M values (TMM) method (Robinson and Oshlack, 2010) on an R framework with the Bioconductor edgeR package (v. 3.28.1) using the default parameters.

### Statistical and bioinformatics analysis

Using Perseus (v.1.6.7.0) (Tyanova et al., 2016), we performed two-sample Welch *t*-tests to identify terminal peptides with a significant alteration upon BB-94 treatment employing a 5% permutation-based FDR filter. Proteolytic events were considered significantly altered if at least one of the N- and C-terminal peptides was significantly altered.

Membrane proteins were annotated with the UniProt Knowledgebase (UniProtKB) keywords “transmembrane” or “GPI-anchor”. Membrane protein topologies and positions of the transmembrane domain and GPI-anchor site were retrieved from UniProtKB, and further analyses were performed using Microsoft Excel.

Other statistics analyses were performed in the R framework (v.3.6.1) with the exactRankTests package (v.0.8-31), psych package (v.1.9.12.31) and pheatmap package (v.1.0.12). Bioinformatics analysis was performed using DAVID (v.6.8) (Huang et al., 2009), STRING (v.11.0) (Szklarczyk et al., 2019), and Cytoscape (v.3.8.0) (Shannon et al., 2003). Sequence logos were created using IceLogo (v.1.0) (Colaert et al., 2009).

### PWM construction and scoring

Flanking sequences surrounding substrate cleavage sites for sheddases were obtained from the MEROPS database (Rawlings et al., 2018) and the previous study (Tucher et al., 2014). Sequences including non-natural amino acids were excluded. PWM construction and PWM score computation were performed essentially as described previously (Imamura et al., 2017). Briefly, the PWM score was computed by summing log_2_ weights for the flanking ±4 residues surrounding each cleavage site.

## Supplemental Information

*Supplemental Figures S1-S7*

**Figure S1. Terminal peptide enrichment by SCX.**

(A, B) Digestion with TrypN yields peptides with at least a +2 charge with Lys or Arg and an α-amino group at the N-terminus (Type C). On the other hand, peptides derived from protein N-termini have a +1 charge with neither Lys nor Arg, and only His-containing peptides with an unmodified N-terminus have a +2 charge (Type A). In addition, digestion with trypsin yields internal peptides with at least a +2 charge (Type B). In this case, C-terminal peptides without His have a +1 charge and those with His have a +2 charge (Type A). Based on these facts, we isolated N-terminal peptides and C-terminal peptides using SCX at low pH, respectively. Note that acetylated N-terminal peptides generated by trypsinILysC digest have a 0 or +1 charge and are isolated with C-terminal peptides.

(C) For N-terminal peptide enrichment, TrypN-digested peptides were dissolved in 2.5% formic acid (FA) containing 30% acetonitrile (ACN) and loaded on the SCX-StageTip. The flow-through fraction was collected. Following TMT-labeling, the isolated N-terminal peptides were analyzed in triplicate.

(D) TMT-labeled peptides were divided into three parts and subjected to C-terminal peptide enrichment using three SCX-StageTips. Peptides were dissolved in 0.15% trifluoroacetic acid (TFA) containing 30% ACN and loaded on the SCX-StageTip. The flow-through and the 0.5% TFA-eluted fractions were separately collected and subjected to single shot analysis.

(E) Efficiency of C-terminal peptide enrichment. As described above, acetylated N-terminal peptides were isolated together with C-terminal peptides.

**Figure S2. Reproducibility of N-terminomics.**

Logarithmized normalized TMT-reporter intensities of N-terminal peptides are plotted against each other for triplicate determinations in each condition, and the corresponding Pearson correlation coefficients are shown.

**Figure S3. Reproducibility of C-terminomics.**

Logarithmized normalized TMT-reporter intensities of C-terminal peptides are plotted for against each other for triplicate determinations in each condition, and the corresponding Pearson correlation coefficients are shown.

**Figure S4. Volcano plots for each cell line in N-terminomics.**

For quantified N-terminal peptides in the respective cell lines, volcano plots were created using Perseus with truncation at the false discovery rate of 0.05 and an artificial within groups variance (S_0_) of 0.1. N-Terminal peptides of membrane proteins are highlighted in color, and proteolytic termini are highlighted with filled circles.

**Figure S5. Volcano plots for each cell line in C-terminomics.**

For quantified C-terminal peptides in the respective cell lines, volcano plots were created using Perseus with truncation at the false discovery rate of 0.05 and an artificial within groups variance (S_0_) of 0.1. C-Terminal peptides of membrane proteins are highlighted in color, and proteolytic termini are highlighted with filled circles.

**Figure S6. Signal peptide cleavage sites and APP γ-cleavage sites, related to Figure 3.**

(A) Distribution of the distance (number of amino acids) from the predicted signal peptide cleavage site to the identified cleavage site.

(B) Sequence logo of the cleavage sites located in the vicinity of the predicted signal peptide cleavage sites (±4 amino acids) generated by iceLogo. The dashed line shows the cleavage site.

(C) Downregulated γ-cleavage sites within the transmembrane domain of APP. The arrowheads indicate the identified cleavage sites, and the numbers indicate the length (number of amino acids) of generated amyloid β peptides.

**Figure S7. PWMs for 16 selected sheddases.**

For the selected sheddases, PWMs were computed based on the sequences of ±4 amino acids flanking the substrate cleavage sites reported in MEROPS.

## Supplemental Data S1-S4

**Table S1-S2. List of all quantified terminal peptides.**

All quantified N-terminal peptides and C-terminal peptides are listed in Table S1 and Table S2 respectively.

**Table S3. List of membrane protein cleavage sites significantly downregulated upon BB-94 treatment.**

**Table S4. Result of PWM score computation.**

